# Development of Molecular Inversion Probes for Soybean Progeny Genomic Selection Genotyping

**DOI:** 10.1101/2022.05.03.490091

**Authors:** Haichuan Wang, Benjamin Campbell, Mary Happ, Samantha McConaughy, Aaron Lorenz, Keenan Amundsen, Qijian Song, Vincent Pantalone, David Hyten

## Abstract

Increasing rate of genetic gain for key agronomic traits through genomic selection requires the development of new molecular methods to run genome-wide single nucleotide polymorphisms (SNPs). The main limitation of current methods is the cost is too high to screen breeding populations. Molecular inversion probes (MIPs) is a targeted genotyping-by-sequencing method that could be used for soybeans that is both cost effective, high-throughput, and provides high data quality to screen breeder’s germplasm for genomic selection. A 1K MIP SNP set was developed for soybean with uniformly distributed markers across the genome. The SNPs were selected to maximize the number of informative markers in germplasm being tested in soybean breeding programs located in the North Central and Mid-South regions of the United States. The 1K SNP MIP set was tested on diverse germplasm and a recombinant inbred line population. Targeted sequencing with MIPs obtained an 85% enrichment for the targeted SNPs. MIP’s genotyping accuracy was 93% overall while homozoygous call accuracy was 98% with less than 10% missing data. The accuracy of MIPs combined with its low per sample cost makes it a powerful tool to enable genomic selection within soybean breeding programs.

## Introduction

Genome-wide genotyping of soybean germplasm lines has been used for many applications, such as quantitative trait loci (QTL) mapping in bi-parental populations, background selection for trait introgression, and genomic selection on progeny rows. These applications typically utilize 100 to 6,000+ molecular markers (Hyten et al., 2004, Kim et al., 2008, Sebastian et al., 2012, Basnet et al., 2022). Since the early 2000s, single nucleotide polymorphism (SNP) markers started to be used in soybean breeding applications (Zhu et al., 2003, Song et al., 2004, Lee et al., 2004). While the marker type has stayed consistent, the technology to genotype SNPs is constantly evolving to decrease the cost of genotyping individual samples with a genome-wide set of molecular markers (Lee et al., 2004, Hyten et al., 2008, Song et al., 2013, Elshire et al., 2011).

One of the first technologies that enabled a large set of molecular markers to be run on soybean germplasm was the GoldenGate assay (Hyten et al., 2008). The development of a Universal SNP Linkage Panel (USLP) 1.0 enabled soybean mapping populations to be genotyped with 1,536 markers within a three-day period (Hyten et al., 2010). The panel was designed so that a high proportion of those 1,536 markers would be polymorphic in any bi-parental mapping population. In bi-parental populations with elite soybean cultivars as parents, the USLP 1.0 had an average of 458 polymorphic markers (informative) segregating in the population (Hyten et al., 2010). This number of polymorphic markers allowed enough coverage in different soybean populations for effective QTL mapping (Kim et al., 2012, Warrington et al., 2015, Phansak et al., 2016). While most markers were monomorphic, the cost savings per sample to obtain the 458 polymorphic markers was cheaper than running each marker individually.

As sequencing cost decreased due to second generation sequencing techniques, methods such as restriction enzyme genotyping-by-sequencing (reGBS) were developed for obtaining genotyping information (Elshire et al., 2011). Restriction enzyme GBS has both advantages and disadvantages when compared to the USLP 1.0. The biggest benefit of reGBS was the significantly decreased cost per sample. This enabled reGBS to be used for multiple genomic applications such as QTL mapping, genomic selection, and diversity analysis (Ravelombola et al., 2021). While the cost of reGBS was cheaper than a genotyping array, data quality became an issue. Restriction enzyme GBS has significant missing data due to low coverage of the same SNPs across samples. To reduce this missing data, imputation methods have been applied to fill in the missing data (Wickland et al., 2017). Meanwhile, the low read coverage of each SNP also causes heterozygous alleles to be under-called in final datasets even after imputation (Nazzicari et al., 2016). Soybean breeding often derives lines in the F_4_ generation which will have on average 12% heterogeneity in their breeding lines. This heterogeneity will likely be under-called using reGBS.

Because reGBS has high rates of missing data and under calls heterozygous alleles, a 6K Illumina Infinium chip (BARCSoySNP6K) is still actively used by the soybean community (Song et al., 2020, Beche et al., 2021, Li et al., 2021). This chip allows researchers to obtain low missing data with high accuracy of calling heterozygotes and has an easy workflow for processing samples and data analysis. Despite the BARCSoySNP6K having a higher cost than reGBS, many soybean researchers and breeders are still willing to use this method for the increase in data quality and simple data analysis.

High quality data with the cost savings of second-generation sequencing can likely be obtained with targeted GBS methods (Teer et al., 2010). Targeted GBS methods can enrich for a set number of small fragments that are known to contain SNPs. This allows for selectively sequencing the enriched fragments to a high read depth that produces a high enough data quality sufficient for breeder’s applications (Mamanova et al., 2010). This high sequencing depth at targeted loci produces enough reads to significantly reduce missing data to the point where imputation is not needed and can accurately identify heterozygous alleles (Teer et al., 2010).

A targeted GBS method that has been successfully applied in multiple species is molecular inversion probes (MIPs) (Niedzicka et al., 2016, Turner et al., 2009, Porreca et al., 2007). Generally, a MIP contains an extension and ligation arm that is complementary to flanking sequence around the targeted SNP. The extension and ligation arms are connected with a backbone sequence that is used for a universal PCR step. Through a hybridization/extension/ligation step, the extension and ligation arm of the probe hybridizes to the targeted sequence. Extension occurs from the extension arm across the targeted sequence. The polymerase used does not have a 5’ to 3’ exonuclease activity, so that extension stops at the ligation arm. Ligation occurs between the extended DNA and the MIP ligation arm to circularized the MIP. Then universal PCR primers complementary to the MIP backbone amplifies the circularized MIP as a linear amplicon. The universal PCR will amplify multiple circularized MIPs that contain the same backbone but targeted different SNPs. For multiple samples, this PCR is used to add dual barcodes for sample identification and the Illumina sequences required for sequencing on Illumina platforms. This method has been demonstrated to be able to target SNPs ranging from a few up to 55,000 in a single MIPs reaction with high accuracy and cost efficiency (Turner et al., 2009, Waalkes et al., 2018).

Genomic selection in progeny rows requires extremely low per sample cost for obtaining informative genome-wide markers in the population being selected upon. These progeny rows will come from many populations with different parental combinations. When the parents of bi-parental populations have been genotyped with a high density of markers then a universal SNP set like the one developed for the GoldenGate USLP 1.0 could be used in genomic selection and be effective across many populations derived from different parents. By developing a universal SNP set of 1000 SNP markers, breeders could use it with a targeted GBS method, such as MIPs, for performing genomic selection in soybean progeny rows. The number of informative markers obtained from a 1000 SNP set will be sufficient to impute the higher density of markers from the parents to the offspring. This would allow breeders to reduce genotyping costs by running the same SNP set on multiple breeding populations. Our objectives were to demonstrate that MIPs could be used as a targeted GBS method for low density SNP genotyping and develop a universal MIPs SNP set that could be used for soybean breeding programs focused on breeding for the North Central and Mid-South regions of the United States.

## Results

### SNP selection for probe set

Multiple methods were explored for the selection of an optimum set of 1,000 SNPs that could make up the MIPs SNP set for genomic selection. The target germplasm for the 1K SNP set is elite breeding lines coming from public soybean breeding programs across the North Central and Mid-South soybean producing regions of the United States. Ideally, this 1K probe set should maximize the number of polymorphic markers in each potential breeding cross and have even coverage across the genome. This will provide enough informative markers for each individual cross without having to create custom marker panels for each population in a breeding program.

Six different methods were tested to determine which method would likely have the most polymorphic markers across breeding programs and have markers distributed evenly across the genome. From the six different methods tested, three methods gave a similar level of polymorphic markers across all target germplasm. The BIN_PIC method produced on average 443 polymorphic markers for a random bi-parental population with elite soybean parents, Song_PIC_MIN_2 produced 445 polymorphic markers and Song_PIC_MIN_0.1 produced 430 polymorphic markers. The other three methods had fewer estimated polymorphic markers with Haploview Tagger having 323 polymorphic markers, Haploview_BIN 312 and Random 314.

While the average number of polymorphic markers is important, the distribution of these markers is also critical for obtaining effective coverage across the genome. The Song_PIC_MIN_2 method has fewer large gaps than the BIN_PIC method when looking at the distribution of markers across the physical space of the genome (Figure 1A, 1B). All the large physical distance gaps from the BIN_PIC method fell within the pericentromeric region. This region is highly heterochromatic and severely represses genetic recombination. The BIN_PIC used bins based on genetic distance to help distribute markers evenly across the genome’s genetic space. This led to fewer markers being clustered in the pericentric region when compared to the Song_PIC_MIN_2 method when looking at the distribution of markers across the genome’s genetic space (Figure 1C, 1D).

**Figure 1.**
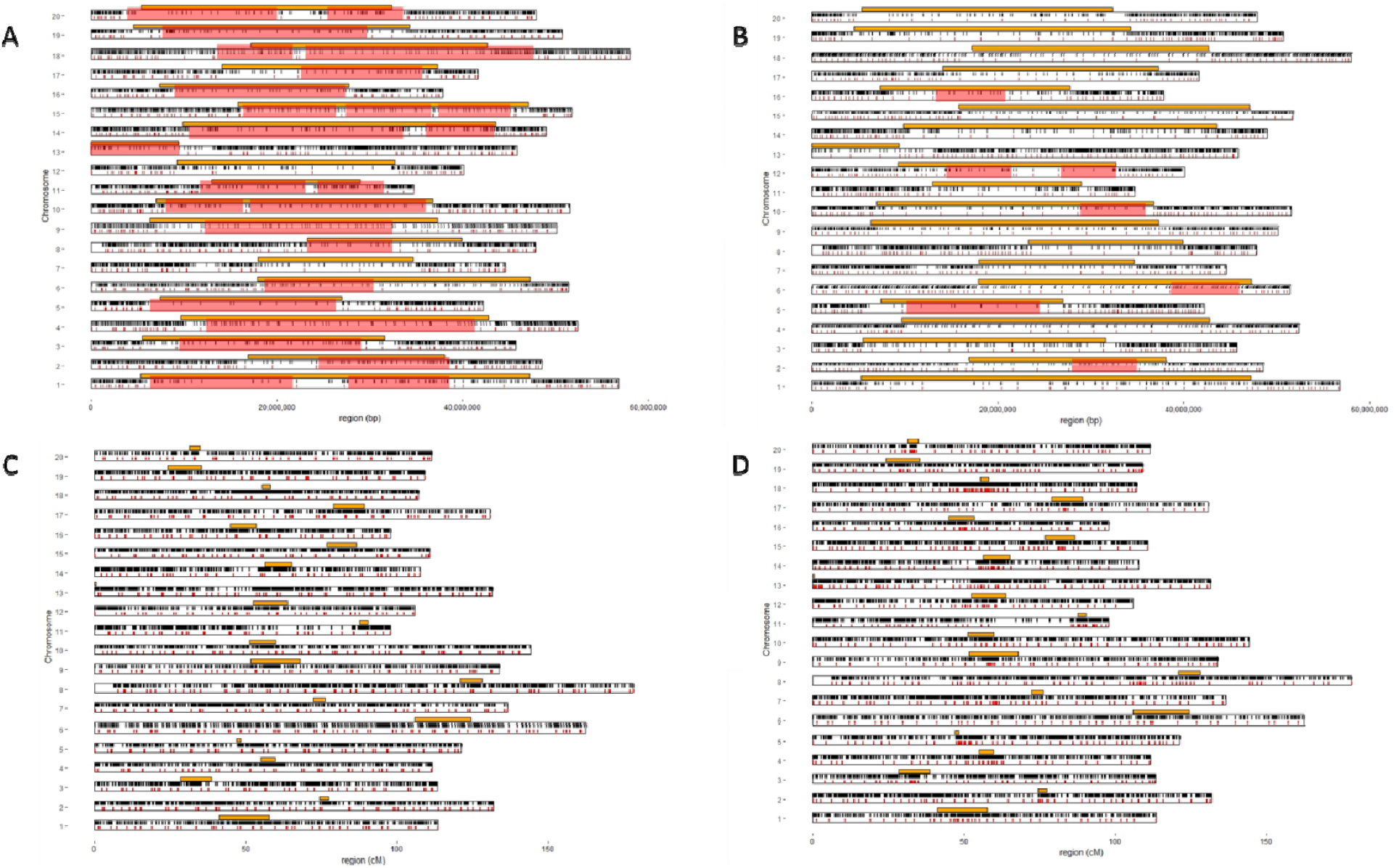
Distribution of MIP 1k selected SNPs verses the BARCSoy6k SNPs. MIP 1k SNPs are indicated with red tick marks, BARCSoy6k SNPs are indicated with black tick marks. Red boxes identify large gaps with no SNPs selected for the 1K MIPs. Orange boxes indicate the soybean pericentromeric regions. A) Distribution of the MIP 1K SNPs selected with the BIN_PIC method across the soybean physical genome. B) Distribution of the MIP 1k SNPs selected with the Song_PIC_MIN_0.2 method across the soybean physical genome. C) Distribution of the MIP 1K SNPs selected with the BIN_PIC method across the soybean genetic distance genome. D) Distribution of the MIP 1k SNPs selected with the Song_PIC_MIN_0.2 method across the soybean genetic distance genome.

The average number of polymorphic markers stayed high for the BIN_PIC method even when the germplasm used to select the markers were split by their respective breeding program (Table 1). The average number of polymorphic markers had a range of 338 to 443 estimated polymorphic markers. The breeding programs with the fewest number of estimated polymorphic markers within populations were Nebraska, Ohio, Southern Illinois and Tennessee. The number of polymorphic markers from the BIN_PIC method was very similar to the two Song methods and considerably outperformed the Haploview and random methods (Table 1).

**Table 1.** Average number of single nucleotide polymorphisms (SNPs) in the 1k MIPs probe set that can be exected to be polymorphic for biparental mattings involved in various combinations for each public soybean breeding program germplasm.

| **Program** | **Number of lines** | **Random** | **Haploview_BIN** | **Haploview_Tagger** | **BIN_PIC** | **Song_PIC_MIN_0.1** | **Song_PIC_MIN_0.2** |
| --- | --- | --- | --- | --- | --- | --- | --- |
| **University of Guelph** | 31 | 293 | 288 | 293 | 383 | 375 | 387 |
| **University of Illinois** | 328 | 293 | 286 | 300 | 421 | 405 | 417 |
| **Iowa State University** | 97 | 280 | 289 | 298 | 421 | 407 | 422 |
| **Kansas State University** | 36 | 253 | 269 | 274 | 386 | 369 | 380 |
| **Michigan State University** | 51 | 293 | 288 | 302 | 412 | 400 | 414 |
| **University of Minnesota** | 354 | 322 | 323 | 332 | 443 | 434 | 448 |
| **University of Missouri** | 51 | 297 | 294 | 305 | 412 | 401 | 410 |
| **University of Nebraska** | 216 | 253 | 244 | 253 | 373 | 357 | 371 |
| **The Ohio State University** | 38 | 242 | 249 | 257 | 351 | 342 | 355 |
| **Purdue University** | 25 | 269 | 269 | 277 | 405 | 387 | 400 |
| **University of Guelph -**<br>**Ridgetown** | 13 | 286 | 308 | 312 | 421 | 409 | 426 |
| **South Dakota State University** | 33 | 292 | 294 | 304 | 409 | 402 | 418 |
| **Southern Illinois University** | 9 | 240 | 244 | 259 | 338 | 335 | 347 |
| **University of Tennessee** | 3 | 240 | 229 | 235 | 352 | 343 | 353 |

### Read alignment of MIPs

The 1K Soy MIP SNP set was used to screen 96 landrace accessions that have previously been screened with the SoySNP50K chip. A total of 26,631895 raw, 150 bp reads were obtained for 93 out of 96 samples. Three samples did not produce enough sequencing data likely due to a low DNA quality. After QC filtering, 25,343,687 reads were kept (95%), of which 22,811,487 reads aligned to the targeted SNP regions. This was equal to an enrichment of 85% for the targeted sequences from the total raw reads.

This enrichment of the targeted sequence led to good read coverage across the targeted SNPs. The average read coverage per SNP in each accession was 245x. A total of 808 probes had an average of 10 reads or more for all the soybean accessions while 53 probes had on average over 1000 reads or more (Figure 2). After eliminating the 192 probes with an average of < 10 reads, SNPs were called on the remaining probes. The concordance of the SNP calls from the 1K data to the SoySNP50K chip was 98% for the 93 PI accessions. The PI accessions are highly inbred with very little heterozygosity. The 98% accuracy obtained from these accessions are reflective of mostly calling homozygous genotypes.

**Figure 2.**
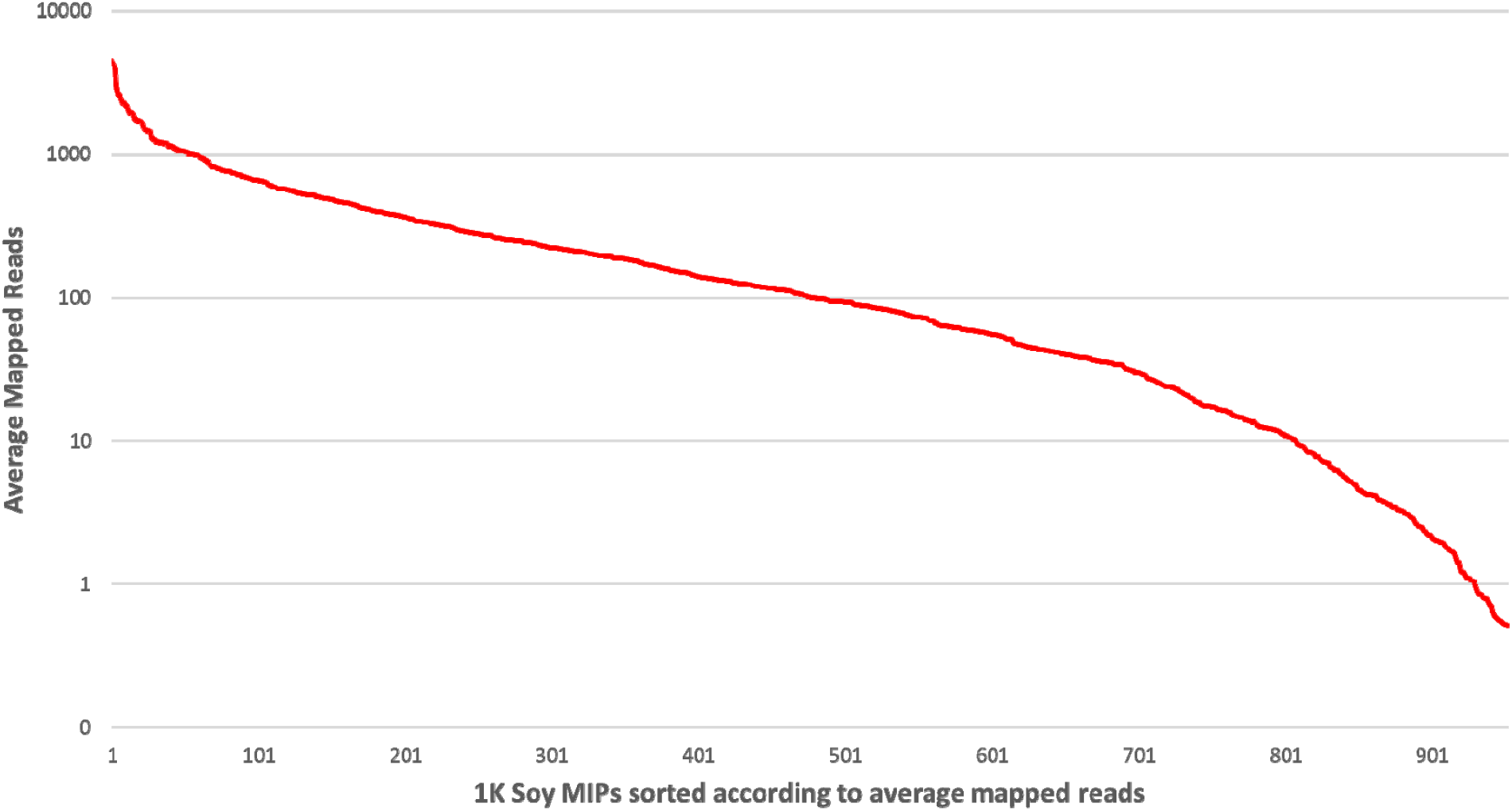
Performance of the MIP markers used to genotype the 96 PI soybean accessions. Standard deviation of MIP probes on the 96 lines represented with error bars.

### Genotype accuracy

To further access genotyping accuracy and the amount of missing data of MIPs genotyping for homozygous and heterozygous genotypes, a RIL population of Essex by Williams82 was genotyped with the 1K MIP SNP set. These RILs have been previously genotyped with the SoySNP50K chip (Song et al., 2016). The SoySNP50K chip includes all the 1K MIP SNPs. The 96 lines were sequenced to an average MIP read depth of 47x (Figure 3). A total of 74 probes aligned to the wrong target and 199 probes had > 30% missing data. These 273 probes were eliminated from the concordance analysis. Six of the genotypes had > 30% missing data. A total of 292 probes out of 725 remaining markers that were polymorphic (40% polymorphism) in the RIL population.

**Figure 3.**
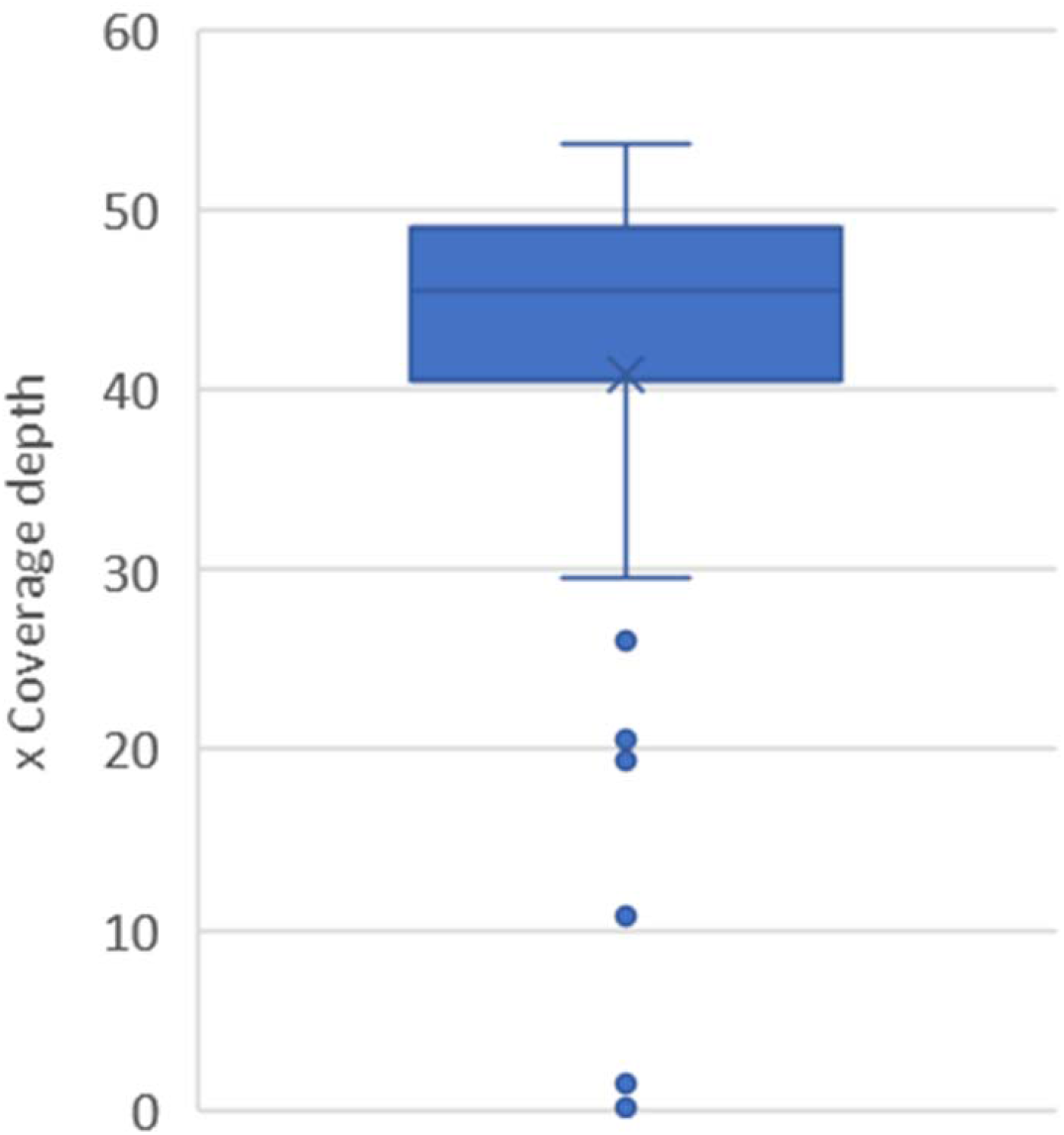
Average sequencing coverage depth for the 1K MIPs genotyping of the Essex x Williams82 recombinant inbred lines.

Concordance of the MIPs genotyping was determined by comparing the SNP genotyping of the RILs performed by Song et al., (2016) to the 1K MIP genotyping. This allowed for determining genotyping accuracy of homozygote and heterozygote SNP genotypes called with MIPs. In addition, multiple SNP calling parameters of GATK were explored to determine their effect on genotyping accuracy (Table 2). Overall, accuracy was above 93% for the genotyping with MIPs. The alignment sensitivity, mapping quality, read depth, and GATK confidence call had limited effect an overall genotyping accuracy and homozygous SNP genotyping accuracy. The different SNP calling parameters did influence heterozygous genotype calls and missing data. The median for heterozygous genotyping accuracy increased to over 90% when the mapping quality parameter of 30 was used. A read depth of 5 vs 10 increased the median accuracy from 90% up to 94%. However, the higher read depth increased the average accuracy for heterozygous calls by only 2% and increased missing data to 13.2%.

**Table 2.** Single nucleotide polymorphism (SNP) calling parameters and affect on accuracy using concordance between MIPs genotyping and the same SNP genotyped with the SoySNP50k Infinium chip on 90 Essex x Williams 82 recombinant inbred lines. MIPs removed with the wrong targets (wrong ref/alt OR concordance < 80%), MIPs with >30% missing data removed, genotypes with > 30% missing data removed

| Parameter | Setting | Overall |  |  | Homozygous |  |  | Heterozygous |  |  | Missing Data |  |  |
| --- | --- | --- | --- | --- | --- | --- | --- | --- | --- | --- | --- | --- | --- |
|  |  | Average | Median | SD | Average | Median | SD | Average | Median | SD | Average | Median | SD |
| Alignment Sensitivity | very sensitive, local | 93.43% | 96.94% | 8.55% | 97.65% | 98.12% | 2.16% | 72.23% | 86.36% | 29.88% | 9.25% | 4.48% | 15.61% |
|  | very sensitive, end to end | 93.64% | 97.37% | 9.02% | 98.35% | 98.82% | 1.74% | 69.49% | 81.82% | 29.20% | 6.93% | 1.37% | 17.19% |
|  | sensitive, local | 93.34% | 96.93% | 8.73% | 97.58% | 98.07% | 2.18% | 72.73% | 89.47% | 30.15% | 9.72% | 4.45% | 16.87% |
|  | sensitive, end to end† | 93.66% | 97.53% | 8.89% | 98.22% | 98.51% | 1.75% | 70.55% | 83.33% | 29.15% | 6.93% | 1.37% | 17.19% |
| Mapping Quality | 10 | 93.64% | 97.53% | 8.91% | 98.22% | 98.51% | 1.75% | 70.49% | 83.33% | 29.21% | 6.93% | 1.37% | 17.19% |
|  | 20 | 93.66% | 97.53% | 8.92% | 98.22% | 98.50% | 1.75% | 70.67% | 83.33% | 29.17% | 6.93% | 1.37% | 17.19% |
|  | 30† | 94.00% | 97.42% | 8.73% | 98.30% | 98.50% | 1.61% | 73.94% | 90.00% | 30.95% | 6.49% | 1.03% | 17.53% |
| Read Depth | 5† | 93.54% | 97.63% | 9.21% | 97.89% | 98.47% | 2.68% | 74.32% | 90.48% | 30.79% | 8.57% | 1.37% | 20.88% |
|  | 10 | 92.37% | 96.84% | 11.19% | 96.39% | 98.05% | 7.00% | 76.14% | 94.12% | 32.23% | 13.20% | 4.11% | 22.66% |
| GATK Confidence Call | 5 | 93.55% | 97.65% | 9.46% | 98.01% | 98.11% | 2.25% | 74.12% | 90.11% | 30.56% | 8.75% | 1.71% | 20.74% |
|  | 10 | 93.72% | 97.67% | 9.16% | 98.07% | 98.77% | 2.70% | 74.31% | 90.48% | 30.77% | 8.82% | 1.71% | 20.79% |
|  | 20† | 93.89% | 97.85% | 9.06% | 98.05% | 98.65% | 2.33% | 74.86% | 90.65% | 30.65% | 8.80% | 1.74% | 20.88% |
| †Final parameters selected for MIPs SNP calling pipeline |  |  |  |  |  |  |  |  |  |  |  |  |  |

## Discussion

We have shown that MIPs can be used to successfully genotype targeted SNPs in soybean. The SNP targeting probes were able to enrich sequence obtained for the selected SNPs by 85%. This high enrichment is higher than the 77.3% enrichment obtained in the non-model organisms smooth and Carpathian newts (Niedzicka et al., 2016). The newts had 80% of the 248 MIPs with a median coverage within one order of each other (Niedzicka et al., 2016). This was a more uniform coverage of MIPs than obtained with the soybean 1K probe set. The main difference in median coverage is likely related to the rebalancing of the MIPs in the newts genotyping. Rebalancing individual probes involves increasing the concentration of the MIPs that produces lower than average read counts and decreasing the concentration of MIPs that produces higher than average read counts. This rebalancing will achieve a more even distribution of average read coverage for each MIP (Niedzicka et al., 2016). When using an unbalanced MIPs set in humans, the uniformity of probe read coverage was similar to the results obtain with the soybean MIPs set (Mamanova et al., 2010). Rebalancing of the starting concentration of individual MIPs can easily be done when the probes are individually synthesized. Through further refinement, the average read coverage for the 1K MIP SNP set can be improved for soybean. This will enable more samples to be multiplexed at a lower average sequencing depth per sequencing run.

The 98% accuracy of genotyping homozygous calls for the soybean 1K MIP SNP set was comparable to the 99% accuracy reported in other species (Niedzicka et al., 2016, Teer et al., 2010). However, the accuracy of calling heterozygotes was lower than expected in this study. One potential reason is the original Illumina genotyping of these RILs were performed with DNA extracted from a single F_5_ plant that derived the RIL. The MIPs genotyping used leaf tissue from a composite of five plants grown from the RIL. The sampling of only five plants from within the line may not be enough to capture all the heterogeneity present within the line. Also, by only sampling five plants there is likely unequal sampling of the two alleles present in a RIL that is heterogenous for a SNP. This could affect the accuracy of calling heterozygous SNPs. Therefore, a deeper sequencing depth or sampling more plants within a line has the potential of increasing accuracy for calling heterozygotes in heterogenous lines when higher accuracy is needed. The overall genotyping accuracy of the inbred lines was greater than 93% which is sufficient for most genomic selection applications.

To achieve the greatest number of informative markers in soybean breeding populations, multiple methods were explored on selecting the 1K MIP SNP set. The Bin_PIC and Song methods effectively have the same number of polymorphic SNPs in all possible bi-parental populations and outperformed both Haploview methods and selecting SNPs randomly. The Bin_PIC and Song methods had on average a 43-44% polymorphic rate while the other three SNP selecting methods had a polymorphic rate of 31-32%. The polymorphic rate with the Bin_PIC and Song methods is higher than the original 1536 USLP 1.0. The 1536 USLP 1.0 had 30% of the markers polymorphic between any two random elite parents (Hyten et al., 2010). This higher polymorphism rate with the MIPs set likely comes from the 1K MIPs SNP set being selected from 50,000 SNP markers, while the original 1536 USLP 1.0 was selected from only 3000 SNPs. This indicates that even though the 1K MIPs SNP set has 35% fewer SNP markers, it will still produce a similar number of polymorphic markers as the 1536 USLP 1.0.

The main difference between the BIN_PIC and the Song methods is how the SNPs are distributed across the chromosomes. By forcing a specific number of SNPs within each bin, the BIN_PIC method ensures that SNPs are distributed evenly across the genetic space of the genome. The Song methods had distinct clustering throughout the genome. This mainly occurs around the percentromeric regions. These percentromeric regions have very high linkage disequilibrium in soybean populations and do not require a high number of markers across its physical region to cover the haplotype variation (Song et al., 2015). The Haploview approach did not perform as well in producing informative polymorphic markers as the Bin_PIC and Song methods. This is likely because Haploview is focused on identifying SNPs that tag the most alleles. This is not necessarily the same thing as capturing the most polymorphic markers and caused the method to underperform in producing a high number of informative markers.

With the high number of informative markers, MIPs has the potential to be run on a large number of breeding lines required to effectively deploy genomic selection or other applications requiring a low density of markers to be run on germplasm. The implementation of dual barcodes for MIPs provides the capability of multiplexing over a thousand samples in a single run. Currently, we have successfully genotyped 1152 samples in a single sequencing run on an Illumina NextSeq 500 with the soybean 1K MIPs SNP set. This both increases throughput and decreases cost per sample. Also, the protocol for MIPs is straightforward and amendable for automation. Using an automatic liquid handler such as the Mantis has enabled the processing of multiple 384 well plates in a day by one person.

MIP’s capability for high sample throughput, high genotyping accuracy, high rate of polymorphic markers in the 1K MIPs SNP set makes it a viable genotyping technology for implementing a low density of markers for genomic selection in soybean breeding populations.

## Materials and Methods

A total of 96 accessions from the USDA germplasm collection and 96 recombinant inbred lines from the cross of Essex by Williams82 (ExW82) were used in this study (Table S1). All lines have been previously genotyped with the SoySNP50K Infinium chip (Song et al., 2016, Song et al., 2015). Leaf tissue was collected from one trifoliate/plant/cultivar for the 96 accessions or 20 leaf discs from five plants/cultivar for the ExW82 RILs. The leaf tissue was lyophilized followed by DNA extraction from the leaf tissues using a CTAB method as described by Happ et al., (2019). DNA samples were quantified with Quantifluor dsDNA system (Promega, Madison, WI, USA) on a Synergy 2 plate reader (BioTek, Winooski, VT, USA) by following manufacturer’s instructions.

SNP markers included in the 1K Soy MIPs SNP set are a subset of the existing BARCSoySNP6K chip (Song et al., 2020). The goal was to select 1000 SNPs that both achieve uniform genome coverage and maximize the number of polymorphic SNPs in any possible bi-parental breeding population formed by crossing elite breeding lines from public soybean breeding programs that participate in the Northern Unform Soybean Tests. Number of polymorphic SNPs was evaluated by using a representative subset of 1285 elite breeding lines entered into USDA Northern Uniform Soybean Tests during the past 10 years. Each breeding line was genotyped with the BARCSoySNP6K. Eighty-four SNPs were removed for having more than 20% missing data or having minor-allele frequency less than 5%. Forty-one SNPs were removed because they did not map to a chromosome in the version 2 genome assembly. A total of 5875 SNPs were remaining for downstream analyses and were candidates for inclusion in the 1K Soy MIPs SNP set. Genetic positions of the SNPs on the BARCSoySNP6K were taken according to the Essex-by-Williams82 (ExW82) genetic map published by Song et al. (2020). The genetic positions of SNPs not positioned on the ExW82 genetic map were interpolated using locally estimated scatterplot smoothing (LOESS) in R relating the physical positions to genetic positions.

The university program and number of lines genotyped per program are displayed in Table 1. All possible bi-parental crosses among all lines, as among all lines within a program were simulated and the number of polymorphic SNPs for each possible cross was enumerated for each method. Six different methods for the selection of SNPs were used and compared: Random, Haploview_BIN, Haploview_Tagger, BIN_PIC, Song_PIC_MIN_0.1, and Song_PIC_MIN_0.2.

### Random

A set of 1000 SNPs was randomly selected from the BARCSoySNP6K. Random sampling of SNPs was repeated 10 times.

### Haploview_BIN

Each chromosome was broken down into bins 5 cM in length for a total of 501 bins. The Haploview software (Barrett et al., 2005) was used to analyze the SNPs in each of the bins and select SNPs that tag the most haplotype alleles. In the end, the number of bins with selected SNPS was 492 with 453 bins having two SNPs and 39 bins only having one SNP.

### Haploview_Tagger

The number of tag markers assigned to each chromosome was based on each chromosome’s genetic length. The program Haploview 4.2 Tagger was used to select the SNPs using an LOD cutoff of 3.0 (Barrett, 2009).

### BIN_PIC

Each chromosome was broken down into bins 5 cM in length for a total of 501 bins. Polymorphism information content (PIC) of each SNP in each of the 14 breeding programs was calculated as PIC = 1 - [(*p*^2^ + *q*^2^) - 2*p*^2^*q*^2^] (Botstein et al., 1980), where *p* is the major allele frequency and *q* is the minor allele frequency. A weighted average PIC score was calculated across all the breeding programs, with the weights being the number of lines within each breeding program (Table 1). The two SNPs in each bin with the highest weighted PIC were selected. Four bins only contained one SNP, for a total of 998 selected SNPs.

### Song_PIC_MIN_0.1

Song et al. (2013) deployed an algorithm to select SNPs that both have high genotyping quality scores and uniformly cover the genome, with different densities for euchromatic and heterochromatic regions.

This method calculated an index score that combined an Illumina design score and “priority score”. An iterative algorithm was developed to select a certain number of SNPs with specified physical distance among them, with the physical distance determined by the ratio of genetic map distance to physical map distance in euchromatic versus heterochromatic regions. The needed density of selected SNPs was determined to be five times greater in euchromatic regions than heterochromatic regions. Gaps between SNPs were determined, and for the smallest gap on any given chromosome a “SNP selection index score” was calculated for the flanking markers of the gap. The SNP with the smallest index score was removed. This procedure was iterated until the desired number of SNPs was selected for each euchromatic and heterochromatic region on each chromosome. Further details of this procedure are provided in Song et al. (2013). This study implemented the same algorithm with the exception that the “SNP selection index score” was replaced by PIC. A minimum PIC value of 0.1 and 0.2 was used for the methods Song_PIC_MIN_0.1 and Song_PIC_MIN_0.2, respectively.

MIPgen (Boyle et al., 2014) was used to design a set of 998 MIPs across the *G. max* genome. The 998 markers (referred to as 1K) are a subset of the BARCSoySNP6K Infinium Chip and the SoySNP50K Infinium Chip (Song et al., 2020, Song et al., 2013). The 998 SNPs were selected using the BIN_PIC method described above. The MIPs were designed to capture a 200-bp sequence with the targeted SNP in the middle of the sequence and filtered based on logistic score. Loess Function in R was used to estimate cM positions for the BARKSoySNP6K positions based on the cM positions from ExW82 population (Song et al., 2016).

Nine hundred and ninety-eight MIPs were individually synthesized by Integrated DNA Technologies (Coralville, IA, USA) at 0.2 nmole/probe (Table S2). The probes were diluted 1:10 with nuclease free water. The probes were then pooled together at equal molar. To phosphorylate the pooled probes a T4 polynucleotide kinase kit (ThermoFisher, Waltham, MA, USA) was used by following the manufacturer’s instructions. In brief, a 20 μl phosphorylation reaction consisted of 15 μl of diluted pooled probes, 1x T4 DNA ligase buffer, 1mM ATP, 0.5U/μl of T4 polynucleotide kinase. The reaction was incubated at 37°C for 30 min, followed by an inactivation of the kinase at 75°C for 10 min.

The hybridization, probe extension, and ligation were carried out in a single step. The 6.34 μl reaction/well was carried out in a 384-well PCR plate. It consisted of individually pipetting 3.34 μl of 12 ng/μl DNA samples (~40 ng genomic DNA) into individual wells of a 384-well PCR plate. The 3 μl master mix consisted of 0.88 μl pooled phosphorylated probes (approximately 1000:1 DNA to probe ratio), 1x Ampligase DNA ligase buffer (Epicentre Technologies, Madison, WI), 0.5 μl of 5M Betain (Sigma Aldrich), 3.25 U AmpliTaq DNA polymerase (ThermoFisher, Waltham, MA, USA), 16 U of DNA Ampligase (Epicentre Technologies, Madison, WI), 0.016 mM dNTP, and 0.08 μl water. The master mix was dispensed into each 384 well using a Mantis Liquid Handler (Formulatrix, Bedford, MA, USA). The reaction was heated at 98°C for 10 min followed by continuous incubation at 60°C for approximately 24 hrs. The heated lid for the PCR machine was set to 105°C for the first 30 minutes and then changed to 80°C after the first 30 minutes of the incubation.

After the hybridization/extension/ligation step, a 2 μl mix of 6.4 U of Exo I (New England Biolabs, Ipswich, MA, USA), 32 U of Exo III (New England Biolabs, Ipswich, MA, USA), 1x Ampligase DNA ligase buffer, and 1.16 μl water was automatically dispensed into each 384-well using a Mantis Liquid Handler (Formulatrix, Bedford, MA, USA) to digest uncircularized MIPs and DNA templates. The reaction was incubated at 37°C for 30 min followed by 95°C for 2 min.

PCR amplification of the circularized MIPs occurred individually for each sample by using 1.25 μl of circularized MIP capture taken from the previous step, 0.5 μl of 10 μM indexed forward and reverse primers (Table S3), and 3 μl of 2x iProof HF Master Mix (BioRad, Hercules, CA, USA). The 3 μl of 2x master mix was dispensed using a Mantis Liquid Handler (Formulatrix, Bedford, MA, USA). The PCR was carried out with the following conditions in a 384-well plate: 98 °C/30 s; 21 cycles of 98°C/10 s, 60°C/30 s, 72°C/30 s; 72°C/60 s, holding at 12 °C.

All PCR products from previous step was pooled by mixing 1μl/sample/well, then the mixed libraries were separated on 1.5% agarose gel for 45 min at 95V along with 50 bp (Fisher, Hampton, NH, USA) and 100 bp DNA ladders (GoldBio, St. Louis, MO, USA). The sharp band at ~286 bp was excised and purified with gel extraction kit (Qiagen, Hilden, Germany) by following manufacturer’s instructions. Alternatively, the pooled PCR products were purified using sbeadex PCR clean-up kits (LGC, Biosearch Technologies) by following the manufacturer’s instructions to purify the PCR band at 286 bp. The purified PCR product was quantified on Qubit (ThermoFisher, Waltham, MA, USA) with Qubit™ RNA HS Assay Kit (ThermoFisher, Waltham, MA, USA) first, then further quantified precisely on QuantStudio 6 plex (ThermoFisher, Waltham, MA, USA) by using the KAPA library quantification kit Illumina platforms (Roche, Basel, Switzerland).

Based on the quantity from qPCR, the purified pooled libraries at 1.5 pM were loaded onto an Illumina NextSeq 500/550 instrument with a Mid Output kit v2.5 (150 cycles) (Illumina, San Diego, CA, USA) using custom sequencing primers (both index and amplicon primers) (Table S4). The sequencing run consisted of a 150 bp single read, 8 bp reads for index 5 and index 7, respectively. The sequencing outputs were processed on The Holland Computing Center, UNL with bcl2fastq2 conversion software (v2.17, Illumina), which combines primary sequencing output (BCL files) from a run and converts/splits them into FASTQ files based on the indexes associated with each sample.

Prior to alignment of reads to the reference sequence, raw reads were filtered with Trimmomatic to remove adapter sequence contamination, truncated reads, and reads with overall low base quality (Bolger et al., 2014). The filtered reads were then aligned to a custom reference genome, which consisted of 200 bp flanking regions around each targeted SNP for each of the 998 SNPs, using Bowtie 2 (Langmead and Salzberg, 2012) in end-to-end mode with the sensitive setting (Niedzicka et al., 2016). After aligning, reads with a mapping quality score of less than 5 were discarded using Samtools. SNP genotypes were then called using the GATK4 tools HaplotypeCaller and GenotypeGVCFs, where calls with a confidence score of less than 20 were removed.

## Acknowledgments

This work was supported by Nebraska Soybean Board project #1723, Development of Next Generation Sequencing Application for Improving Soybean and the North Central Soybean Research Program project: Increasing the Rate of Genetic Gain for Yield in Soybean Breeding Programs. This work was completed utilizing the Holland Computing Center of the University of Nebraska, which receives support from the Nebraska Research Initiative. This material is based upon work supported by the National Science Foundation Postdoctoral Research Fellowship in Biology (Grant No. IOS-1710790). Any opinions, findings, and conclusions or recommendations expressed in this material are those of the author(s) and do not necessarily reflect the views of the National Science Foundation. The authors acknowledge the Minnesota Supercomputing Institute (MSI) at the University of Minnesota for providing resources that contributed to the research results reported within this paper. URL: http://www.msi.umn.edu

## Supporting Information

Table S1. List of Soybean accessions used in this study.

Table S2. The SNP name, alleles, location and sequence of the molecular inversion probes contained within the 1K Soy MIP SNP set.

Table S3. MIPs amplification and indexing primers. Dual indexing combinations are used to multiplex large number of samples in one sequencing run.

Table S4. Custom sequencing primers used for sequencing MIP amplified probes

## Notes

### Competing Interest Statement

The authors have declared no competing interest.

